# Mice deficient in nucleoporin Nup210 develop peripheral T cell alterations

**DOI:** 10.1101/330522

**Authors:** Annemarie van Nieuwenhuijze, Bert Malengier-Devlies, Bénédicte Cauwe, Stephanie Humblet-Baron, Adrian Liston

**Author notes:** Co-last, co-corresponding authors: Adrian Liston,; Stephanie Humblet-Baron.

## Abstract

The nucleopore is an essential structure of the eukaryotic cell, regulating passage between the nucleus and cytoplasm. While individual functions of core nucleopore proteins have been identified, the role of other components, such as Nup210, are poorly defined. Here, through the use of an unbiased ENU mutagenesis screen for mutations effecting the peripheral T cell compartment, we identified a Nup210 mutation in a mouse strain with altered CD4 / CD8 T cell ratios. Through the generation of Nup210 knockout mice we identified Nup210 as having a T cell-intrinsic function in the peripheral homeostasis of T cells. Remarkably, despite the deep evolutionary conservation of this key nucleopore complex member, no other major phenotypes developed, with viable and healthy knockout mice. These results identify Nup210 as an important nucleopore complex component for peripheral T cells, and raise further questions of why this nucleopore component shows deep evolutionary conservation despite seemingly redundant functions in most cell types.

## Introduction

An understanding of the genetic requirements for T cell development has been built upon the analysis of murine and human T cell immunodeficiencies. These studies have identified genes that have roles in differentiation, function, maintenance or homeostasis of T cells, with mutation leading to loss of the T cell population [1]. While most mutations leading to T cell-deficiency have clear lineage-specific functions, there is a fascinating class of mutations in genes that are widely expressed and have basic cell-biological functions, such as gene regulation (DNMT3β, SP110 [2; 3]), chromatin remodelling (SMARCAL1 [4]), and metabolism (adenosine deaminase, nucleoside phosphorylase [5; 6]. While complete loss of many of these genes would be anticipated to result in embryonic lethality, based on critical functions in cell biology, identified mutations tend to have T cell-specific defects. This observation is thought to be a result of selection bias, where only those point mutations mild enough to retain sufficient function for most cells result in viable offspring. It is not clear why T cells are sensitive to mild mutations that other cells can tolerate, however this may be related to the rapid rate of proliferation of the early stages of T cell differentiation [7; 8]. Regardless, in multiple cases T cells have functioned as the “canary in the coal mine”, acting as a phenotyping read-out for mild mutations in critical cell biology genes.

Nup210 (or gp210) was the first nucleopore-associated protein to be discovered, and was initially thought to promote the fusion of inner and outer nuclear membranes during nucleopore assembly [9; 10]. However, Nup210 is not ubiquitously expressed in all tissues, and the analysis of nucleopore complex (NPC) composition during mouse embryogenesis and in naturally Nup210-deficient cell lines showed that Nup210 is dispensable for the assembly or stability of the nucleopore complex (NPC) [11; 12; 13; 14]. While this result may discard Nup210 as an essential component of the NPC, its symmetrical localisation as a membrane ring around the nuclear pore, which is also observed for the yeast homologue Pom152 [15], and high conservation across eukaryotes, was suggestive of an important function in cell biology. More recently, it was shown that shRNA knockdown of Nup210 in myoblasts and embryonic stem cells induced apoptosis and completely abrogated their differentiation into myotubes and neuroprogenitor cells [16]. Further studies have suggested that Nup210 is acting as a scaffolding protein for transcriptional complexes such as Mef2C, and that the tissue-specific expression is most likely a driver for the specialisation of NPCs in different cell types, thereby playing a role in the regulation of cell fate [17]. This cellular function has been studied in most detail in myocyte culture [17], however the role of NUP210 at the organism level has not been studied.

The dual identity of T cells as both critical coordinators of the immune system and also highly sensitive indicators of disturbed cellular processes makes them attractive targets for unbiased genome-wide genetic screens. Here we used an ENU mutagenesis screen for altered peripheral T cell phenotypes to identify an I476T point mutation in Nup210 which skews the CD4:CD8 compartment ratio. Generation of Nup210 knockout mice validated a T cell-intrinsic function for Nup210. The surprising viability of Nup210 knockout mice leads to the perplexing question of the driver of deep evolutionary conservation of a seemingly largely redundant nuclear pore factor.

## Results

### Mutation in conserved nucleoporin Nup210 alters composition of the T cell compartment

As part of an unbiased screen for genetic control of the peripheral T cell compartment, ENU-exposed C57BL/6 mice were bred to *Foxp3*^*GFP*^ females to generate a standard F2 intercross pedigree [18]. The resulting ENU mutants were screened for altered ratios of CD4 and CD8 T cells in the peripheral blood. Within one pedigree, individuals were identified with a decreased CD4:CD8 ratio in the peripheral blood. Intercrossing of affected individuals resulted in a mutant strain which consistently demonstrated a decreased CD4:CD8 ration in the spleen at 5-6 weeks of age (**Fig. 1a**). All-exon sequencing of affected individuals identified an A→G nucleotide substitution at nucleotide 1469 of the Nup210 gene, which was confirmed by Sanger sequencing (**Fig. 1b**). This mutation in exon 11 (**Fig. 1c**) resulted in a predicted isoleucine to threonine change at amino acid 476 (Nup210^I476T^). The mutation was located in an invariant amino acid in a region of Nup210 highly conserved throughout vertebrates (**Fig. 1d**).

**Figure 1:**
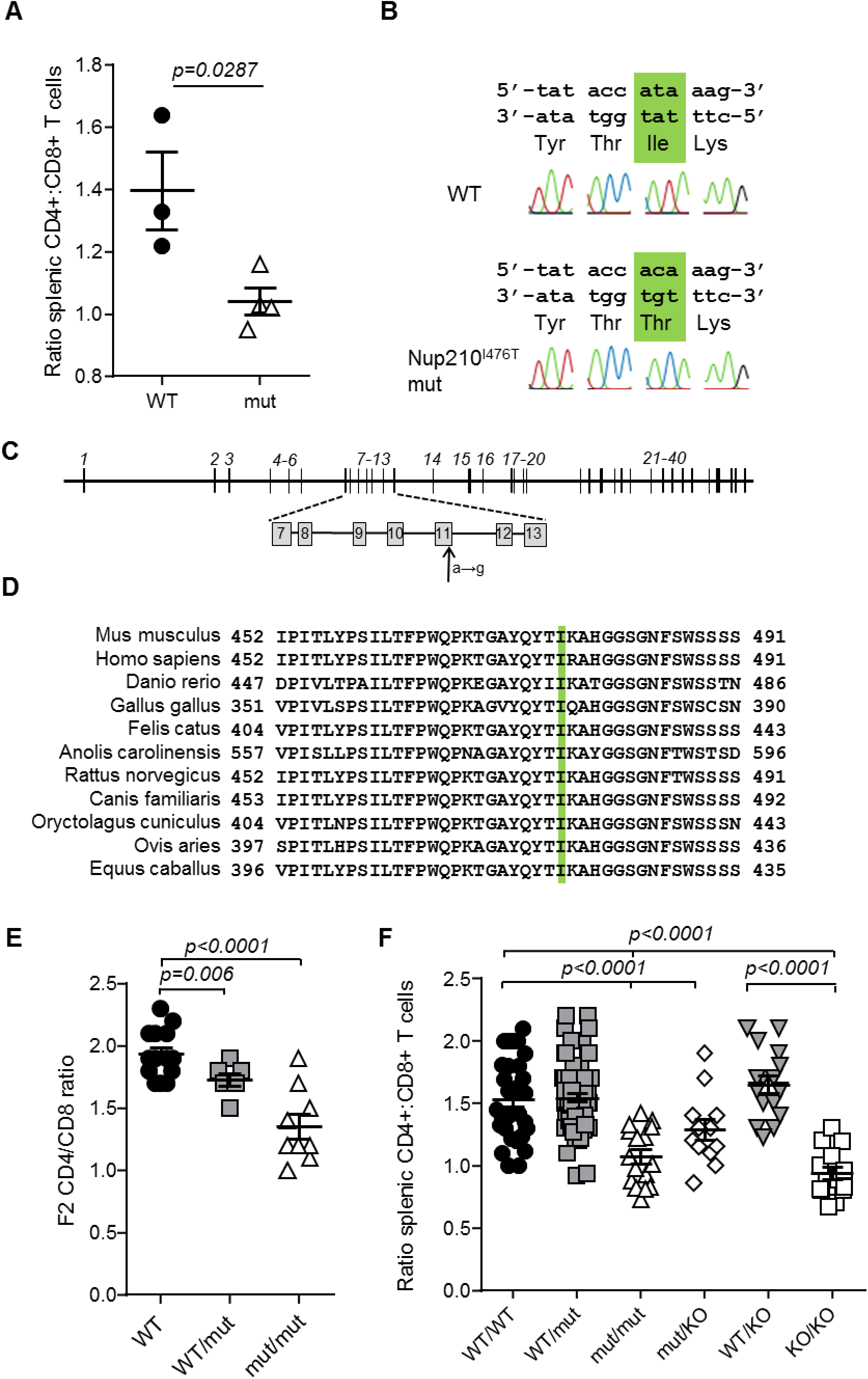
Altered ratio of peripheral CD4:CD8 T cells in Nup210 mutant mice. ENU mutagenesis generated a *Nup210*^*I476T*^ mouse strain, identified by peripheral blood screening for T cell composition. **(A)** Peripheral blood from 5-6 week old wildtype (WT) and Nup210^I^476^T^ mutant (mut) mice was assessed for the CD4:CD8 T cell ratio by flow cytometry (n=3,4). **(B)** Sanger sequencing of *Nup210* in WT and Nup210^I^476^T^ mutant mice confirmed an A to G mutation, resulting in an isoleucine to threonine change at amino acid 476. **(C)** Schematic overview of the 40 exons of the *Nup210* gene, including the location of the I476T mutation in exon 11 (arrow). **(D)** Conservation of the mutation site between the mouse, human, zebrafish, chicken, cat, lizard, rat, dog, rabbit, sheep and horse homologous sequences. **(E)** Confirmation of germline transmission of the *Nup210*^*I476T*^ mutation in F2 offspring of the *Nup210*^*I476T*^ mutant founder mouse and resulting splenic phenotype (n=14,7,9). **(F)** Replication of the *Nup210*^*I476T*^ mutant phenotype (spleen) in a complementation cross (n=33,71,16,16,20,17). Mean±SEM, with individual biological replicates.

Nup210 is thought to be a key component of the nuclear pore complex (NPC), forming a membrane ring around the NPC [15]. Due to the lack of known biology linking Nup210 to T cell-specific processes, we sought to validate the mutation through an F2 phenotyping cross and a complementation cross. First, *Nup210*^*I476T*^ mice were crossed with wildtype mice to produce an F1 generation and then intercrossed to produce an F2 generation. Genotyping for the *Nup210*^*I476T*^ allele allowed identification of wildtype, heterozygous and homozygous mice, while other, unlinked, ENU-induced alleles would be randomly segregated across the pedigree. Phenotyping of the F2 cross found a replication of the original finding, a reduction in the CD4:CD8 ratio in *Nup210*^*I476T*^ homozygous mice (**Fig. 1e**). Second, for a complementation cross, mice bearing a Nup210KO allele were generated using EUCOMM ES cells, with an insertion of a LacZ/Neo cassette between exons 3 and 5 (**Fig. S1**). Mice bearing one copy of the *Nup210*^*I476T*^ allele and one copy of the *Nup210*^*KO*^ allele phenocopied the *Nup210*^*I476T*^ homozygous mice (**Fig. 1f**). Together, these data validated Nup210 as a novel genetic control mechanism controlling CD4:CD8 ratio.

### Nup210 knockout mice are viable and manifest a subtle T cell-specific phenotype

While no known functions of Nup210 are linked specifically to T cell biology, the rapid rate of proliferation of early T cells is known to sensitise this lineage to minor genetic insults in general cell biology components, such as the chromatin condensing unit kleisin beta [7]. Based on the essential function of Nup210 in the nuclear pore complex, and the lethal phenotype that results from knock-down of Nup210 in cell lines [19], it was assumed that the *Nup210*^*I476T*^ allele was a mild hypomorph, with sufficient function maintained to prevent cell death in lineages beyond early T cell stages. The generation of mice bearing the *Nup210*^*KO*^ allele allowed direct testing of this hypothesis, by intercrossing *Nup210*^*het*^ mice to create *Nup210*^*KO*^ mice. Surprisingly, intercross of *Nup210*^*het*^ mice produced wildtype:*Nup210*^*het*^:*Nup210*^*KO*^ mice at the Mendelian 1:2:1 ratio. *Nup210*^*KO*^ mice were viable, demonstrated no visual abnormalities or altered total bodyweight or the weight of key immunological organs (**Fig. 2a**). Histological screening of the organs was unremarkable (data not shown). Complete knock-out of Nup210 in *Nup210*^*KO*^ mice was confirmed by Western blot (**Fig. 2b**), leaving the perplexing finding that an evolutionarily conserved component of the nuclear pore complex is largely redundant for life.

**Figure 2.**
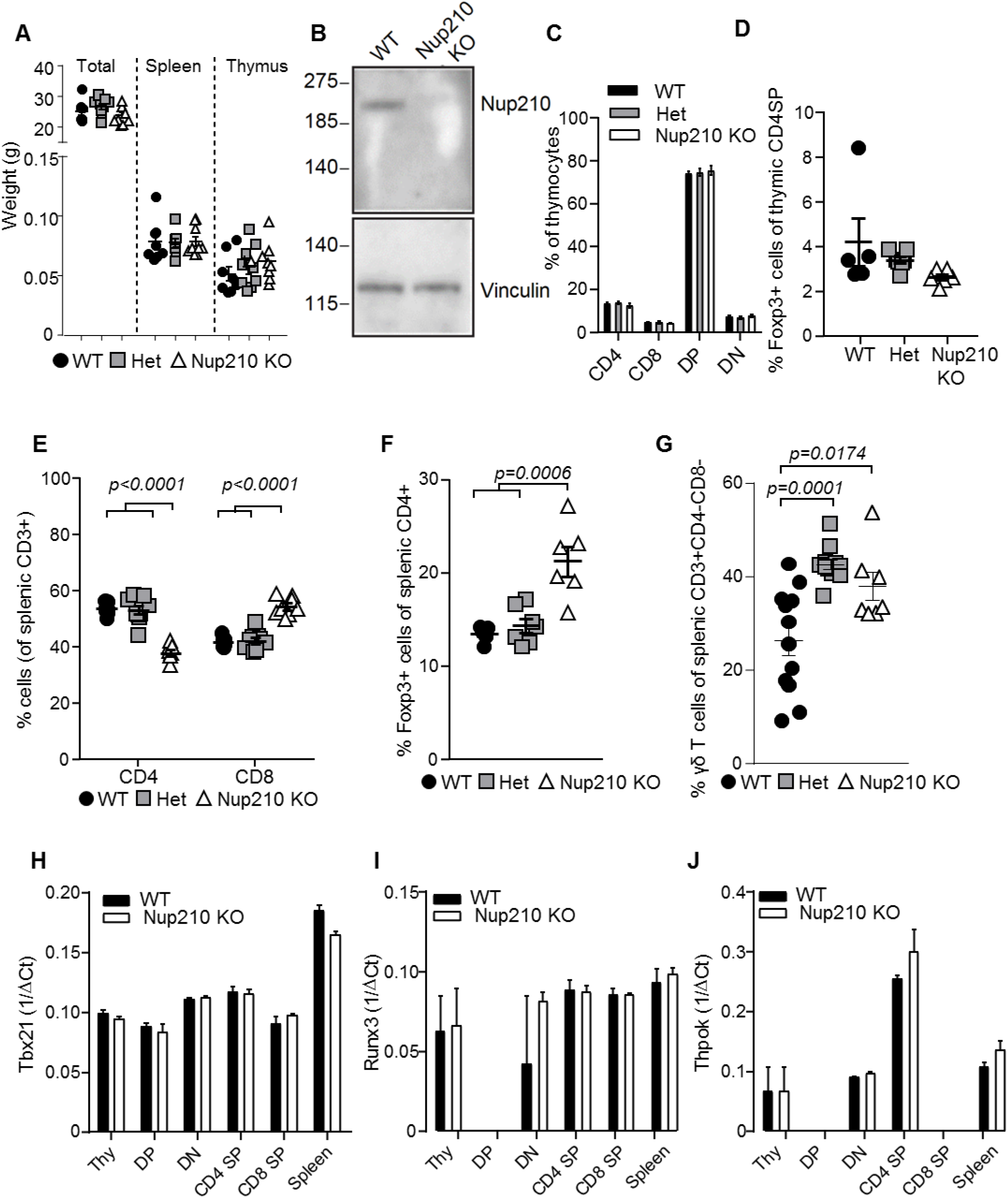
Nup210 knockout mice are viable and manifest a subtle T cell-specific phenotype. **(A)** 6 to 11 week old wildtype, *Nup210*^*het*^, *Nup210*^*KO*^ mice were analysed for body weight, spleen weight and thymus weight (n=8,9,9). **(B)** Western blot for Nup210 in the lysates of wildtype and *Nup210*^*KO*^ thymocytes. Vinculin was used to control for protein input (representative of 2 independent experiments). **(C-K)** Wildtype, *Nup210*^*het*^, *Nup210*^*KO*^ mice (n=5,7,6) were assessed by flow cytometry for **(C)** double negative (DN), double positive (DP) and CD4 and CD8 single positive subsets in the thymus; **(D)** Foxp3+ regulatory T cells in the thymic CD4 compartment; **(E)** CD4 and CD8 T cells in the spleen; **(F)** Foxp3+ regulatory T cells in the spleen; and **(G)** γd T cells in the spleen. Mean±SEM, with individual biological replicates. **(H-J)** Wildtype and *Nup210*^*KO*^ mice were used as donors for whole thymocytes, DN, DP, CD4 SP, CD8 SP, and whole splenocytes. Normalised expression of *Tbx21, Runx3* and *Thpok* in each population, through qPCR (n=2 mice per genotype; representative result from n=3 experiments). Mean±SEM.

Early stage T cells are highly susceptible to cellular stress, owing to the rapid proliferation at the double negative (DN) stage of development in the thymus, followed by the precarious double positive (DP) stage, at which “death by neglect” is the default outcome [1]. We therefore tested whether the peripheral T cell phenotype observed in *Nup210*^*KO*^ mice is due to thymic defects. No difference was observed between wildtype and knockout mice in the DN or DP stages, or in single positive (SP) thymocytes which had diverged into either the CD4 or CD8 lineage (**Fig. 2c**), or Foxp3+ regulatory sublineage (**Fig. 2d**). Despite this, the same mice demonstrated the peripheral T cell effects in the spleen, with reduced CD4 T cells, increased CD8 T cells (**Fig. 2e**), increased Foxp3+ regulatory T cells (**Fig. 2f**) and increased γδ T cells (**Fig. 2g**). Key transcription factors known to impact the CD4-CD8 lineage decision in the thymus, Tbx21, Runx3 and Thpok, remained unchanged throughout thymic differentiation (**Fig. 2h-j**). Together, these results demonstrate that the role of Nup210 in shaping the peripheral T cell compartment is a novel, peripheral-specific function, rather than a reflection of thymic sensitivity to basic cellular processes.

Further investigation into the function of Nup210 in T cells was led by compartment analysis. Mixed bone-marrow chimeras were set up, where wildtype and *Nup210*^*KO*^ haematopoietic stem cells reconstitute an irradiated mouse. This approach allows the competitive comparison of wildtype and *Nup210*^*KO*^ T cells in a context where the host environment is directly shared, thus excluding any effects of non-haematopoietic origin. The CD4:CD8 ratio disturbance still developed in *Nup210*^*KO*^ cells in this context (**Fig. 3a**), demonstrating that the function of Nup210 in driving this phenotype is intrinsic to T cells. The same result was also demonstrated when reconstituting Rag-deficient mice, where contamination from residual host-derived T cells can be excluded (**Fig. 3b**). The demonstration that the function of Nup210 was T cell-intrinsic led to further fine phenotyping of *Nup210*^*KO*^ mice. Compared to heterozygous littermates, *Nup210*^*KO*^ mice manifested a Th1-bias, with increased numbers of IFNγ-producing CD4 and CD8 cells, while Th2 and Th17 cells were unchanged (**Fig. 4a,b**). Proliferation rates of both CD4 and CD8 cells were increased (**Fig. 4c**), while the CD69 activation marker was also increased on CD4 T cells (**Fig. 4d**). Together, these data were suggestive of a pro-inflammatory status of Nup210-deficient mice. An immunological challenge to induce inflammation, using the collagen-induced arthritis model, did not, however, identify any susceptibility to inflammatory disease (**Fig. S2**). Together, these results demonstrate that Nup210 restrains IFNγ production, however the deficiency effect remains subclinical even under inflammatory conditions, perhaps due to a complementary increase in regulatory T cells (**Fig. 2f**).

**Figure 3:**
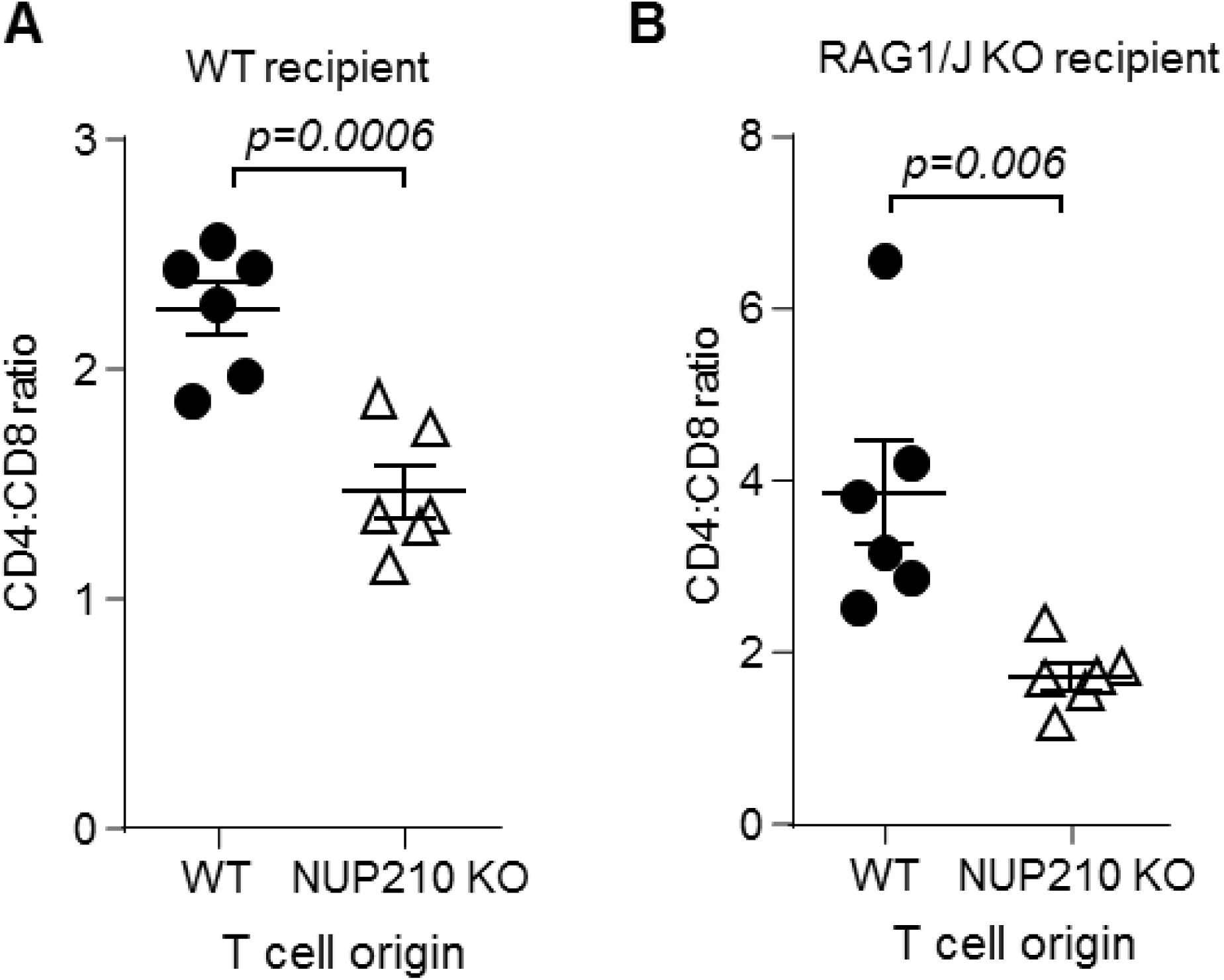
Altered ratio of peripheral CD4:CD8 T cells in *Nup210*^*KO*^ mice is due to functions intrinsic to T cells. **(A)** Wildtype CD45.1 mice were irradiated and reconstituted with 50% bone-marrow from wildtype CD45.1 mice and 50% bone marrow from *Nup210*^*KO*^ CD45.2 mice (n=6). At 8 weeks post-reconstitution, the CD4:CD8 ratio in the spleen was assessed by flow cytometry in both the wildtype (CD45.1+) and *Nup210*^*KO*^ (CD45.2+) compartment. **(B)** *Rag1*^*KO*^ CD45.2 mice were irradiated and reconstituted with 50% bone-marrow from wildtype CD45.1 mice and 50% bone marrow from *Nup210*^*KO*^ CD45.2 mice (n=6). At 8 weeks post-reconstitution, the CD4:CD8 ratio in the spleen was assessed by flow cytometry in both the wildtype (CD45.1+) and *Nup210*^*KO*^ (CD45.2+) compartment. Mean±SEM, with individual data points.

**Figure 4:**
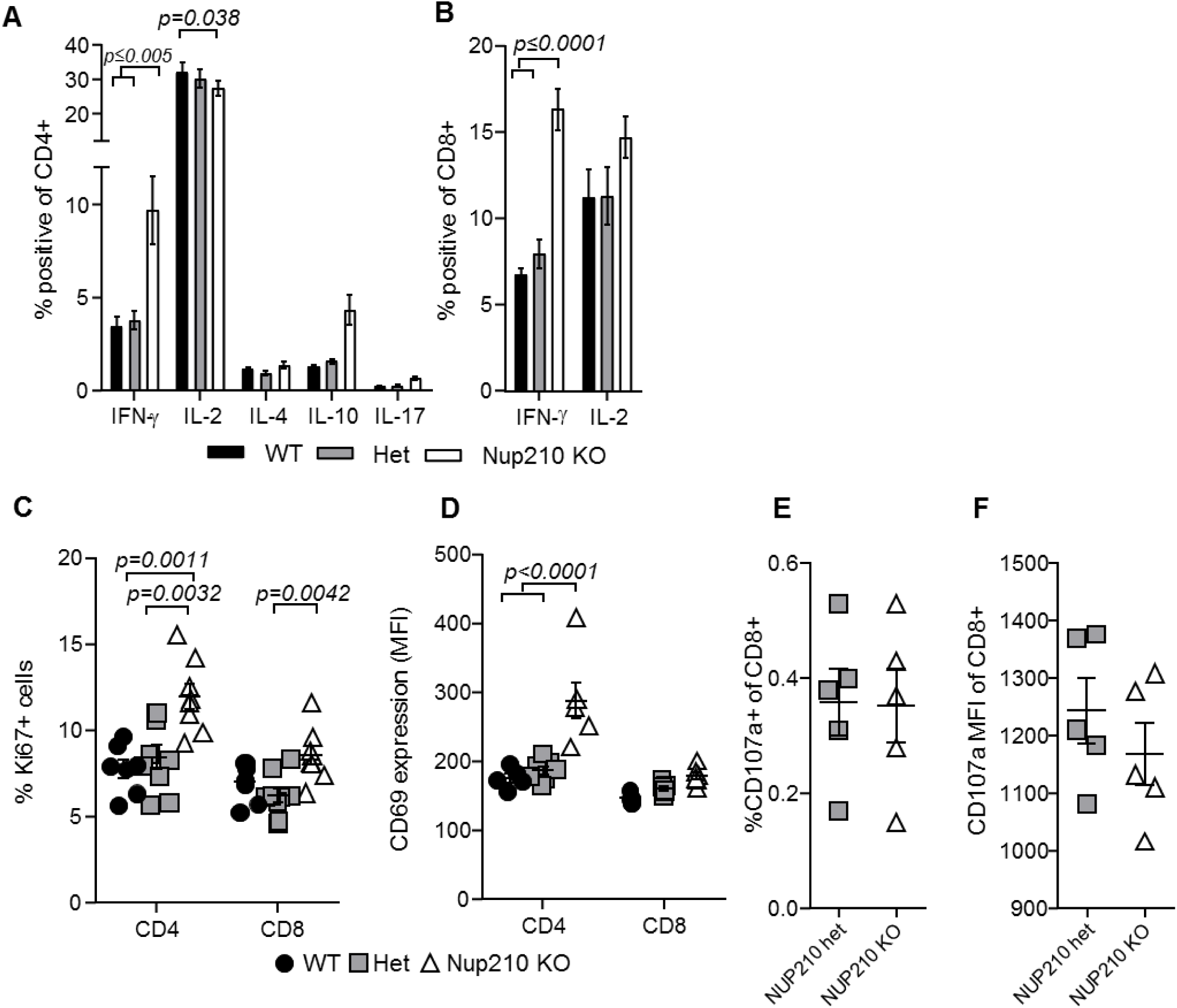
*Nup210*^*KO*^ mice exhibit an inflammatory T cell signature. Splenocytes from wildtype, *Nup210*^*het*^, *Nup210*^*KO*^ mice (n=6,7,5) were assessed by flow cytometry. **(A)** Cytokine production by CD4+ cells or **(B)** CD8+ cells after PMA/ionomycin stimulation. **(C)** Proliferation of CD4+ and CD8+ T cells. **(D)** CD69 expression in splenic T cells. Mean±SEM.

## Discussion

Both the Nup210 mutant mouse strain and the Nup210 knockout mouse strain manifested a disturbance in the peripheral T cells compartment, namely in the ratio of CD4 to CD8 T cells. The observation of general cell biology defects manifesting with T cell components is reoccurring [1], and may lie in the extraordinary rate of proliferating of early stage T cell differentiation in the thymus. Such a model would be consistent with the lethal phenotype observed with Nup210 deficiency in HeLa cells, embryonic *C. elegans* and differentiation embryonic stem cells [19; 20]. Indeed, one of the few known functions of Nup210 is the nucleocytoplasmic transport of the mitosis promoting factor (MPF) [21], and it may play a role in the breakdown of the nuclear envelope during mitosis [22; 23]. Despite the attractive synergy of such a model, analysis of the thymus suggested normal T cell development, with no alterations in the differentiation stages and no alteration in expression of ThPOK, involved in the CD4 linage commitment or Runx3, involved in the CD8 linage commitment [24; 25]. The T cell phenotype, shown here to be T cell intrinsic, instead appears to manifest entirely at the naïve peripheral T cell stage, a relatively quiescent low-activity cell type. This observation, and the observation that cytokine production in activated T cells was altered in Nup210-deficient mice, suggests that the function of Nup210 may lie in gene regulation, such as through altered gene expression [21; 26], rather than in basic cell biology functions such as proliferation. The mechanism by which Nup210 could alter gene expression without entering the nucleus remains unknown [16; 17]; the simplest explanation, altered trafficking through the nucleopore, is not supported by experimental testing [26]. Here Ptprf was an attractive target, with downregulation observed in post-mitotic myotubes following Nup210 depletion [16], and a known function in T cell biology [27]. While this candidate was not observed to be differentially expressed in Nup210-deficient T cells, disturbed gene expression remains an attractive hypothesis for Nup210 function, if only because no obvious alternatives have been proposed. The functional role of Nup210 in T cell activation identified here makes the highly specific link between anti-Nup210 autoantibodies and primary biliary cirrhosis [28], a T cell-mediated autoimmune disease [29], even more intriguing.

The most intriguing, and perplexing, finding from the data in this study is the “negative result” of a weak phenotype in Nup210-deficient mice. Indeed, the very finding that Nup210-deficient mice can be generated is highly surprising. Nup210 is a highly conserved component of the nuclear pore, a complex essential for eukaryotic life owing to the function in trafficking between the nucleus and cytoplasm. It is feasible, indeed, even likely in light of the current results, that Nup210 is not an obligate component of this complex. Compensation by other nuclear pore components is one potential explanation, however it was not observed at the RNA level (data not shown). Regardless, the deep evolutionary conservation of this protein is difficult to explain without a critical non-redundant function in same aspect of cell biology. Indeed, deficiency in Nup210 in HeLa cells and reduction of Nup210 by RNAi in C.*elegans* resulted in greatly reduced viability and early lethality [19], supporting a critical function for this protein. We document here a function for Nup210 in T cells, however the phenotype induced by deficiency is unlikely to explain the evolutionary conservation, especially across species that do not have T cells. One potential explanation is that Nup210 has a critical function which is not revealed under homeostatic conditions, but which nevertheless confers a key survival advantage under certain stress conditions. Indeed, shRNA-mediated depletion of Nup210 in vitro resulted in the upregulation of endoplasmic reticulum stress-specific caspase cascades [30]. Here we tested autoimmune stress (in the context of arthritis induction) and metabolic stress (glucose tolerance after exposure to a high fat diet; data not shown), with no clear phenotype shown, however an infinite range of stress contexts is possible. The generation of these mice opens up the capacity for future exploration of the hidden functions of Nup210.

## Materials and methods

### Mice

All animal experiments were approved by the Animal Ethics Committee of the KU Leuven and performed in accordance with the approved protocol. To generate the *Nup210*^*ENUI476T*^ strain, founder C57BL/6 male mice were treated with 100 mg/kg ENU and bred to *Foxp3GFP* females [31]. First-generation (F1) male offspring were bred back to WT females to produce the second-generation (F2) offspring, which were in turn inter-crossed to produce the third generation for phenotypic screening. Phenotypic screening involved flow-cytometric analysis for CD4 and CD8 in the blood. *NUP210*^*KO*^ mice were generated through the European Conditional Mouse Mutagenesis (EUCOMM) program, by insertion of a LacZ/Neo cassette between exons 3 and 5 (**Fig. S1**).

### Flow cytometry

Surface staining was performed in RPMI containing 2% fetal bovine serum and anti-mouse CD16/CD32 Fc block (from hybridoma supernatant generated in-house). The following antibodies were used in this study: CD3 (145-2C11), CD4 (GK1.5 and RM4-5), CD8α (53-6.7), Foxp3 (FJK-16s), IFNγ (XMG1.2), IL-17 (TC11-18H10), IL-4 (BVD6-24G2), IL-2 (JES6-5H4), IL-10 (JES5-16E3), Ly5.1 (A20), Ly5.2 (104) (eBioscience, CA, USA). Nuclear staining for Foxp3 was performed according to the manufacturer’s recommendations (eBioScience, CA, USA). Intracellular cytokine staining was performed after a 4-hour stimulation with 50ng/mL phorbol 12-myristate 13-acetate and 0.5μg/mL ionomycin (Sigma) in the presence of GolgiStop (Monensin A, BD Biosciences, NJ, USA), using Cytofix/Cytoperm (BD Biosciences, NJ, USA). Dead cells were excluded from analysis by staining with Zombie dyes (BioLegend, San Diego, CA, USA) according to the manufacturer’s instructions. Samples were analysed using a BD FACSCanto II instrument (BD Biosciences) and FlowJo software (Treestar Inc., OR, USA).

### Collagen-induced arthritis

CIA was induced in 6-10 week old mice as previously described [32]. A total of 100μL of chick type II collagen (CII, final concentration 1mg/ml; Sigma, MO, USA) emulsified in complete Freund’s adjuvant containing 5 mg/ml heat-killed *M. tuberculosis* H37RA (BD Difco, NJ, USA) was injected intradermally in two sites at the base of the tail. The injections were repeated 21 days later. Animals were monitored three times weekly for erythema and swelling of limbs, and a clinical score (0–3) was given for each paw. Serum was collected using serum separator tubes (Greiner, Vilvoorde, Belgium) and analyzed for anti-chick collagen type II IgG antibodies by ELISA as described [33]. Standard curves were constructed from pooled sera of CII hyper-immunized DBA/1 mice, set at 100000 units/ml.

### Bone marrow chimeras

Red cell-depleted bone marrow from donor mice was depleted for mature T cells by incubation with biotinylated antibodies to CD3, CD4 and CD8 (eBioscience, San Diego, USA) followed by streptavidin-coupled Dynabeads and magnetic separation according to the manufacturer’s instructions (ThermoFisher, Gent, Belgium). Bone-marrow chimeras were generated by lethal irradiation (9.5 Gy) of recipients, followed by intravenous (i.v.) injection of 5×10^6^ donor cells in saline. Reconstitution was analysed after 8 weeks by flow cytometry.

### Expression analysis

Quantitative RT-PCR was performed on purified mRNA (Trizol reagent, Ambion, Belgium) from cell populations sorted by flowcytometry from thymi of 6-week old mice, using the GoScript Reverse Transcription kit (Promega, Wisconsin, USA) and FastSYBR green reagents (ThermoFisher, Belgium). Control mRNA included mRNA purified from thymocytes and splenocytes. Primer pairs used were as follows: Tbx21 fwd: AGGGGGCTTCCAACAATG; Tbx21 rev: AGACGTGTGTGTTAGAAGCACTG; Runx3 fwd: ACCACGAGCCACTTCAGCAG; Runx3 rev: CGATGGTGTGGCGCTGTA; Thpok fwd: ATGGGATTCCAATCAGGTCA; Thpok rev: TTCTTCCTACACCCTGTGCC; Ppia fwd: GAGCTGTTTGCAGACAAAGTTC; Ppia rev: CCCTGGCACATGAATCCTGG.

Western Blot was performed on thymocytes lysed by sonication in lysis buffer (200 mM NaCl, 50 mM Tris pH 7.5, 1% Triton X-100, 2 mM dithiothreitol, 1 mM EDTA, protease inhibitor (ThermoFisher, Gent, Belgium)). Lysates (20ug) were run on 8% NuPAGE BisTris gels and blotted to a polyvinylidene fluoride transfer membrane using the NuPage electrophoresis system (ThermoFisher, Gent, Belgium) according to the manufacturer’s recommendations. After washing in NCP (147 mM NaCl, 40 mM Tris pH 8, 0.01% Tween), the membrane was blocked overnight at 4°C with 5% non-fat milk in NCP 0.01% Tween. Primary antibodies against Nup210 (Abcam, Cambridge, UK, ab15600, 1:500) or control vinculin (Sigma-Aldrich, St.Louis, USA, V9131, 1/2000), were incubated in NCP 0.01% Tween, 1% non-fat milk. The membrane was washed in NCP 0.01% Tween and the primary antibody was detected with horseradish peroxidase-conjugated anti-rabbit secondary antibody (ThermoFisher, Gent, Belgium, 1:40,000) for Nup210 and HRP-conjugated anti-mouse secondary antibody (Merck Millipore, Darmstadt, Germany, 1:10,000). After washing in NCP 0.01% Tween, blots were developed using the Amersham ECL Prime Western Blotting Detection Reagent (GE Healthcare, Buckinghamshire, UK). The Spectra multicolour high range protein ladder (ThermoFisher, Gent, Belgium) was used to determine the molecular weights of the visualized bands.

## Acknowledgements

The authors wish to thank Denise Klatt and Samuel Ribeiro-Viseu for help with experiments, and Susann Schönefeldt and Jeason Haughton for help with mouse breeding, and the Leuven University Animalium staff for animal husbandry. This work was supported by the VIB and the Belgian Science Policy Office Interuniversity Attraction Poles program (T-TIME). AvN was supported by the Leuven University F+ fellowship, BC was supported by the Fonds Wetenschappelijk Onderzoek (FWO) Flanders.

---

**Supplementary Figure 1:**
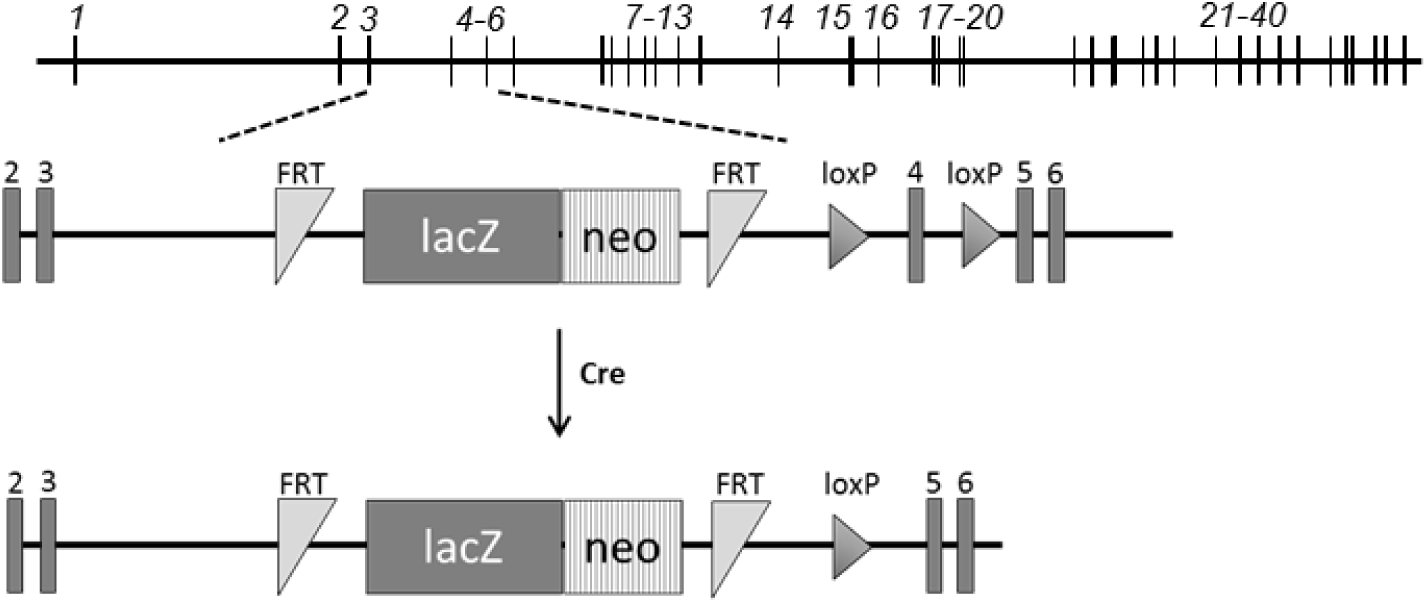
Generation of Nup210-deficient mice. Schematic depiction of *Nup210* locus design in original EUCOMM ES cells.

**Supplementary Figure 2:**
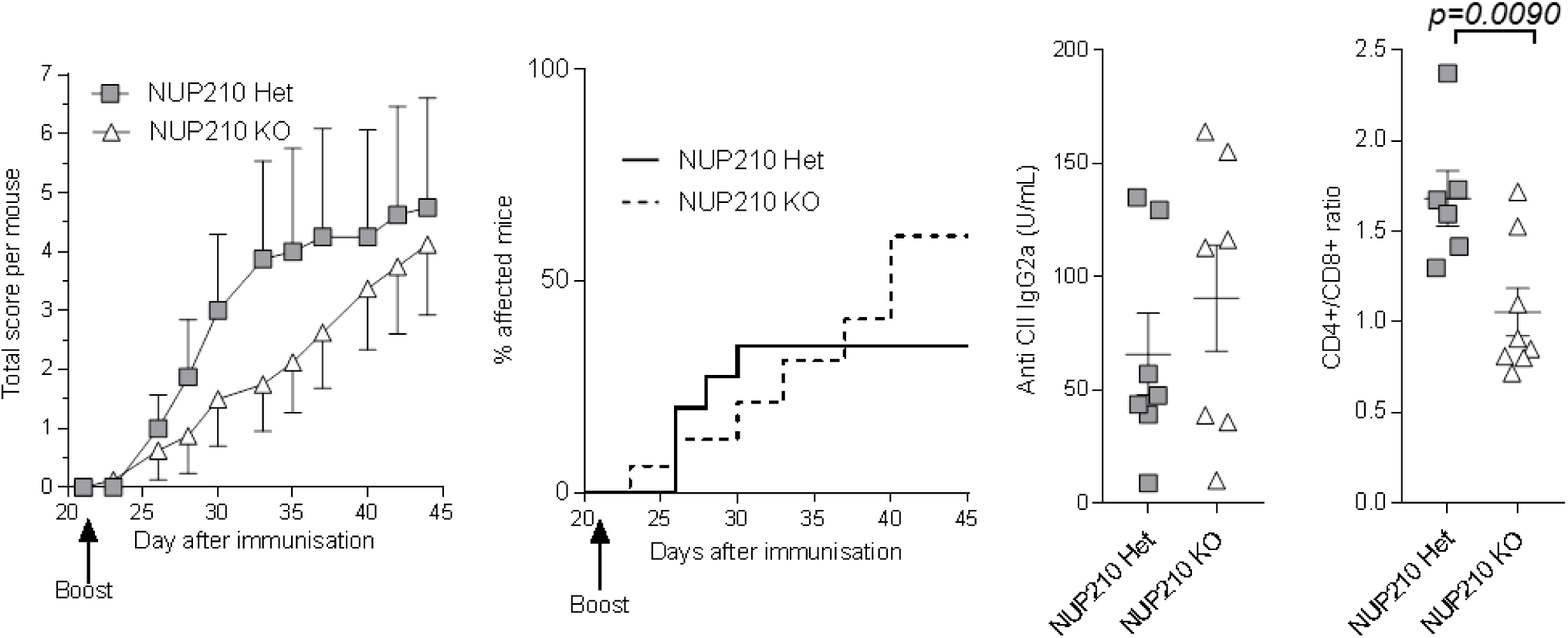
Normal susceptibility to collagen-induced arthritis in Nup210 knockout mice. *Nup210*^*het*^ and *Nup210*^*KO*^ mice were immunised with chick type II collagen (CII) on day 0 and boosted on day 21. Disease incidence and severity was monitored 3 times weekly up to day 45. (n=8,6). **(A)** Total disease score per mouse (maximum 3 per paw). No significant difference by T test comparing the area under the curve calculated for individual mice. **(B)** Incidence of disease (not significantly different, Log-rank Mantel-Cox test). **(C)** Anti-CII IgG2a levels in the serum on day 45 (not significantly different). **(D)** CD4:CD8 T cell ratio in the spleen on day 45, as measured by flow cytometry. Mean±SEM, with individual data points.

